# Making Pastoralists Count: Geospatial Methods for the Health Surveillance of Nomadic Populations

**DOI:** 10.1101/572685

**Authors:** Hannah Wild, Luke Glowacki, Stace Maples, Iván Mejía-Guevara, Amy Krystosik, Matthew H. Bonds, Abiy Hiruy, A. Desiree LaBeaud, Michele Barry

## Abstract

Nomadic pastoralists are among the world’s hardest-to-reach and least-served populations. Pastoralist communities are difficult to capture in household surveys due to factors including their high degree of mobility over remote terrain, fluid domestic arrangements, and cultural barriers. Most surveys utilize census-based sampling frames which do not accurately capture the demographic and health parameters of nomadic populations. As a result, pastoralists are “invisible” in population data such as the Demographic and Health Surveys (DHS). By combining remote sensing and geospatial analysis, we developed a sampling strategy designed to capture the current distribution of nomadic populations.

We then implemented this sampling frame to survey a population of mobile pastoralists in southwest Ethiopia, focusing on maternal and child health (MCH) indicators. Using standardized instruments from DHS questionnaires, we draw comparisons with regional and national data finding disparities with DHS data in core MCH indicators including vaccination coverage, skilled birth attendance, and nutritional status. Our field validation demonstrates that this method is a logistically feasible alternative to conventional sampling frames and may be used at the population level. Geospatial sampling methods provide cost-affordable and logistically feasible strategies for sampling mobile populations, a crucial first step towards reaching these groups with health services.

## Introduction

Nomadic pastoralists defy many basic premises of household demographic surveys, including the assumption that individuals are attached to a geographically stable household, and that this household represents a fixed domestic unit.^1^ By contrast, pastoralist settlements are often highly mobile, moving over large areas of remote terrain with the herds of livestock on which they subsist.^2^ Their domestic arrangements are similarly fluid, as family members reside in different geographic locations to manage these livestock. Combined with the dispersed distribution of their encampments and cultural barriers, these mobility patterns make surveying nomadic pastoralists notoriously difficult.

Despite being among the most underserved populations in Sub-Saharan Africa,^3^ nomadic pastoralists are also underrepresented in the population data used to plan health interventions.^4^ Large-scale household surveys such as the Demographic and Health Surveys (DHS) Program typically use census-based sampling frames, which magnify and institutionalize the issue of mobile pastoralists’ under-enumeration in the original census. Such strategies result in the “statistical invisibility” of nomadic populations.^1^ A lack of accurate data prevents anything but speculative estimates of the global population of mobile pastoralists, but estimates range from 50 to 217.5 million.^4, 5^ Pastoralists globally face threats to their health and livelihood including ecologic disruptions, large-scale development projects, conflict, and protracted humanitarian crises. With mounting concerns about emerging zoonotic pandemic disease, it is critical to find ways of including nomadic populations in household surveys and health surveillance systems.

Numerous methodological approaches to achieving representative samples of mobile populations have been tested, including a “waterpoint approach” in which data is collected at waterholes;^6^ a capture-recapture transect approach similar to that used to monitor wildlife;^7^ random geographic cluster sampling;^8^ and the use of mobile phones.^9^ Many of the approaches explored in these studies have been limited by logistical obstacles or do not lend themselves to being scaled and integrated into data collection exercises such as the DHS.

Geospatial methods present new opportunities for sampling mobile pastoralists, and have been employed with sedentary populations^10, 11^ as well as at the single-village level.^12^ However, geospatial methods have not been used to develop and implement sampling frames at a scale that could be integrated with existing national health surveys. In this study, we develop and assess the utility and feasibility of a geospatial sampling frame to conduct household surveys among a population of mobile pastoralists in a remote region of southwest Ethiopia.

## Materials and Methods

### Study Site and Population

The Nyangatom are nomadic pastoralists numbering approximately 25,000 individuals.^13^ They inhabit a remote lowland area of approximately 2,600 square km on the border between Ethiopia’s Lower Omo Valley and South Sudan. Their settlement pattern consists of mobile camps ranging from several families to two hundred individuals in size, as well as semi-permanent villages associated with areas of seasonal crop cultivation. The composition of all encampment types is fluid, and individuals move between settlements.^14^ There is one market town in Nyangatom in which a health clinic is located. Distances to the clinic from Nyangatom settlements vary greatly by region and season, but often require significant travel by foot. Our target population included women of reproductive age and their under-five-year-old children. We surveyed 342 mothers of reproductive age and recorded data on the health status of 826 of their children 15 years of age and younger, 547 of whom were under five.

### Ethical Precautions

Verbal informed consent from adults and oral assent from minors was obtained from all study participants. Participants’ personal health information (PHI) was protected on researchers’ fully encrypted devices with procedures for de-identification of data during analysis, and no PHI was linked to geospatial data in the public domain. Ethical approval for this study was obtained from the Institutional Review Board for Human Subjects Research at Stanford University School of Medicine, the Ministry of Science and Technology, and the Southern Nations, Nationalities, and Peoples’ Regional Health Bureau in Hawassa, Ethiopia.

### Sampling Design

We used 0.5-meter resolution satellite imagery of the study area in visible and infrared imagery bands, collected within four months of survey administration (WorldView-3, April 26th, 2017). The imagery was first prepared using the Geographic Data Abstraction Library (GDAL) for pan-sharpening, band-reduction, and export of False-Color Infrared images in RGB format, which we used to assist in assessing settlement inhabitance status. The resulting imagery was then converted into an Esri Tile Package and deployed to the ArcGIS Online platform in order to identify settlements. Using the Hanna Immersive Visualization Environment (HIVE) at Stanford’s Institute for Computational & Mathematical Engineering (ICME) consisting of 35 HD screens with an effective resolution of 13440×5400, we tagged actively inhabited settlements and recorded WGS84 geospatial coordinates for use in clustering. We defined settlements as the area within 500 meters of an enclosed camp. Encampments have a distinctive visual profile consisting of an outer ring of kraals and a central collection of huts as well as shade structures, storage platforms, and meeting areas (See Figure 1). Among other populations the composition of settlements will vary, and different features should be used for identification. Since nomadic dwellings are typically constructed from natural materials such as saplings and grass, inhabited versus abandoned settlements can be difficult to distinguish from satellite imagery. Due to the presence of grazing livestock, inhabited settlements contain little interior vegetation while uninhabited settlements tend to be overgrown. We used the presence of significant vegetation in the interior of a settlement as an indicator that the settlement was likely to be uninhabited.

**Figure 1.**
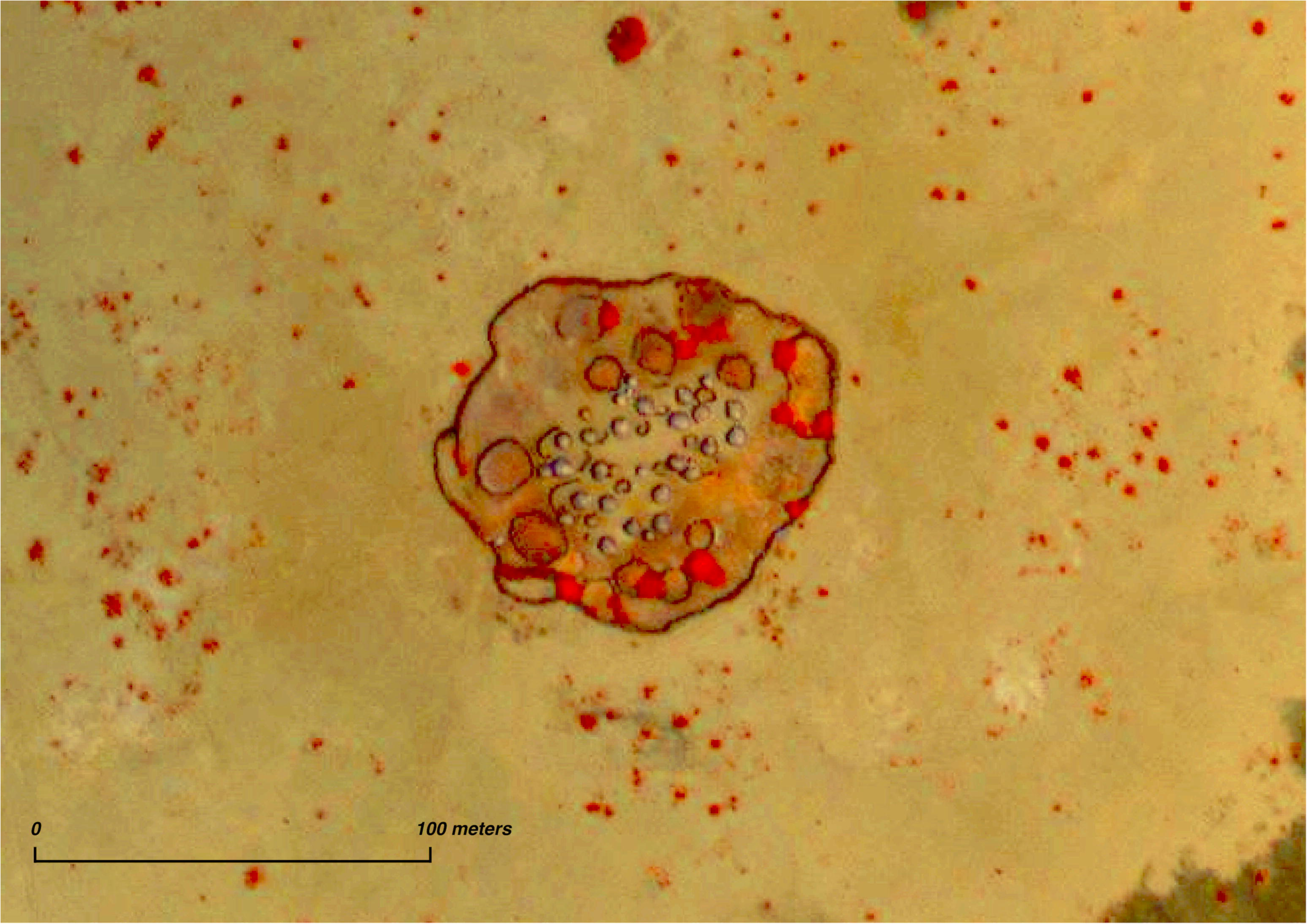
Nyangatom settlement (diameter approx. 100m) with huts, livestock kraals, and perimeter fence viewed using Infrared and Visible Multispectral Imagery (red indicates vegetation.) Captured from imagery covering total land area of 5000km^2^ at a resolution of 0.53m. (Imagery provided by DigitalGlobe Foundation)

As the only available sampling frame of primary sampling units (PSUs) was based on census data and therefore not representative of the target population, we relied on spatial data to divide the entire area into smaller adjacent subregions as homogeneous as possible in terms of population size. We visually identified 225 settlements in the study area, comprising 3099 huts. With ArcMap’s Grouping Analysis Tool, we used settlement elevation and K-nearest neighbors for initial grouping analysis, which produced 15 seed points (the maximum number allowed by the program). We then grouped settlements into 15 clusters with an average of 206 huts per cluster around these points based on elevation and spatial proximity. In order to develop primary sampling units (PSUs), we generated Thiessen polygons from these clusters using the spatial median of settlements within each cluster, thereby minimizing the total distance to all points. This process was reiterated until no further change to cluster membership occurred, producing 15 PSUs covering the entire survey area. Although other approaches such as treating individual settlements as the PSU were considered, we favored clustering since the sparse distribution of settlements would have otherwise made conducting the field survey infeasible. Our approach ensured that the survey would be operationalizable, minimizing travel time within clusters and maximizing field team safety,

We then selected three PSUs to survey using probability proportional to size sampling (PPS) based on the number of huts per settlement as identified in the geospatial data. Although a greater number of sampling units could have been selected to provide an even more representative sample, our decision to sample three clusters balanced reducing sampling variability with the financial and logistical constraints of the field survey (see SI section Sample Size Calculation & Sampling Weights).

Lastly, we carried out a field survey in the selected PSUs. We used individual settlements as identified in satellite imagery as our secondary sampling unit (SSU). We generated a randomized list of all settlements within each cluster, visiting a total of 25 settlements in order to achieve a sufficient sample size (see SI section Sample Size Calculation). Within each settlement we conducted in-person interviews with all women of reproductive age who reported having a birth at any time during the five years preceding the survey. We also collected anthropometric data for the three youngest children aged 0-15 for each woman in our sample. Our survey instruments were developed using questionnaires from the most recent Ethiopian Demographic and Health Survey (EDHS 2016).^15^

### Field Procedures

For each settlement selected for the field survey, researchers used GPS coordinates obtained from satellite imagery to locate nomadic encampments in the field. If upon reaching the settlement it was abandoned or could not be located, we selected the spatially nearest replacement. If no spatially proximate alternate could be identified, we selected the next randomly-assigned numerical ID in the enumeration list as a replacement. All eligible respondents present within a 500m^2^ area surrounding the central point of an enclosed camp tagged from satellite imagery were included. Study personnel adhered to DHS protocols while conducting interviews and obtaining anthropometric measurements. We measured height (cm) and weight (kg) for eligible under-five-year-old children using adjustable ShorrBoard measuring boards and digital SECA scales. Standing height was measured for children aged two to five years, and recumbent length was measured for children less than two years of age.

### Analysis

We conducted descriptive statistical analyses of basic demographic and health indicators for mothers and their three youngest children below the age of 16. As a benchmark for assessing the health status of our sample in comparison with populations from other regions of Ethiopia, we used the 2016 Ethiopian Demographic and Health Survey (EDHS) survey^15^. The 2016 EDHS contains information on 16,650 households and 10,335 rural women. Where EDHS data are available at the regional level, we compare our results to data for the Southern Nations, Nationalities, and People’s Region (SNNPR), of which the Nyangatom administrative district *(woreda)* is a part. Where EDHS data is disaggregated only by urban versus rural residence at the national level, we compare our results to data from rural areas.

For mothers in our sample we report marriage indicators including marital status, number of co-wives, and wife order in polygynous marriages. We calculated fertility indicators based on the number of live births and miscarriages per woman, as well as data on subsequent deaths of children. We also collected data on MCH indicators including antenatal care (ANC) visits, services delivered during the ANC visit, location of child birth, vaccination coverage, and autonomy of women to seek health services (full questionnaire in SI Appendix). Consistent with the DHS program, we collected information on treatment-seeking behavior for symptoms of diarrhea, fever, and cough among under-five-year-old children.

We imputed the month of birth for 284 (52%) under-five-year-old children for whom age was reported in years but not month by using multiple imputation (MI), since age in months was required for the assessment of anthropometric analysis (see SI section Multiple Imputation). Raw height and weight measures were converted to age-and sex-specific z-scores represented by standard deviation (SD) units by using the WHO child growth standards.^16^ We constructed three indicators of anthropometric failure, defined using z-scores less than −2 SD of the median Height-for-Age (HAZ) for stunting, Weight-for-Age (WAZ) for underweight, and Weight-for-Height (WHZ) for wasting. Severe anthropometric failure was defined on the basis of z-scores less than −3 SD of the respective anthropometric scores. Mid-upper arm circumference (MUAC) was also measured (cm) for children aged 6-59 months and used as an alternative assessment to identify Severe Acute Malnutrition (SAM), defined for infants/children with a MUAC<11.5cm^16^ (see SI Methods).

## Results

We interviewed 342 women reporting having given birth during the five years preceding the survey. Maternal age was not recorded because the Nyangatom do not keep track of their own age; however, we assumed a reproductive range of 15-49 years.

### Antenatal Care (ANC) and Delivery

256 of 342 women (75.9%) reported receiving ANC at least once during their most recent pregnancy, 71.6% (95% CI: 56.0-83.3%) of whom visited a skilled provider as defined by the EDHS including nurses, midwives, health officers, health extension workers, and other medically trained personnel. These rates are comparable to those reported by the 2016 EDHS for women regionally in SNNPR, 69.3% of whom received ANC services at least once from a skilled provider. Nyangatom women were 4.0 (3.0-6.0) months pregnant (median, IQR) at the time of their first visit to an ANC provider and made an average of 2.8 (2.4-3.2) visits during their most recent pregnancy. The majority of women in our sample who had given birth to a child under five years of age at the time of the study did so at their home or another home (91.2%, 95% CI: 61.3-98.6%), and in a health facility or clinic in only 6.8% (1.0-36.3%) of all deliveries. In comparison, the 2016 EDHS found that 72.5% of women in the region delivered at home, and in a health facility or clinic for 25.5% of all deliveries. See SI Table 2 for comprehensive descriptive statistics of antenatal care seeking behavior and antenatal services received.

**Table 1.** Marriage, fertility, and mortality indicators of women and their children (Nyangatom, 2017)

| Indicator | Ethiopia DHS 2016 | Nyangatom 2017 |  | p-value<sup>6</sup><br>(1) vs. (2) |
| --- | --- | --- | --- | --- |
|  | Weighted % / mean<br>(1) | n | Weighted % /<br>mean (sd)<br>(2) |  |
| Number of all women of<br>childbearing age interviewed |  | 347 |  |  |
| Marital Status |  |  |  |  |
| Unmarried<sup>1</sup> |  | 1 | 0.2 |  |
| Married / Living together |  | 338 | 97.4 |  |
| Widowed |  | 8 | 2.3 |  |
| Total |  | 347 | 100.0 |  |
| Mean number of wives per husband<br>(if married) |  | 338 | 1.8 (1.0) |  |
| Wife Order (if other wives) |  |  |  |  |
| 1 |  | 71 | 39.8 |  |
| 2 |  | 69 | 38.7 |  |
| 3 |  | 23 | 12.7 |  |
| 4+ |  | 11 | 5.6 |  |
| Missing / Refused |  | 7 | 3.1 |  |
| Total |  | 181 | 100.0 |  |
| Ownership of farm animals<sup>2</sup> | 87.6 | 328 | 96.8 | 0.02 |
| Fertility, children ever born and<br>living<sup>3</sup> |  |  |  |  |
| Mean number of pregnancies per<br>woman of childbearing age |  |  | 5.6 (3.4) |  |
| Mean number of miscarriages per woman of childbearing age |  |  | 0.6 (0.9) |  |
| Mean number of live births per woman of childbearing age (sd)<sup>4</sup> | 4.2 (2.5) |  | 5.0 (3.1) | 0.10 |
| Mortality |  |  |  |  |
| Mean number of dead children per woman of childbearing age |  |  | 0.8 (1.2) |  |
| Male child deaths (mean) |  |  | 0.5 (0.8) |  |
| Female child deaths (mean) |  |  | 0.3 (0.7) |  |
| Mean number of living children per woman of childbearing age (sd)<sup>5</sup> | 3.7 (2.2) |  | 4.2 (2.4) | 0.13 |

**Source:** Nyangatom data.
<sup>1</sup> Includes 'Never married' and 'Divorced/separated' in DHS 2016.
<sup>2</sup> In DHS, numbers are from rural areas and include: cows, bulls, other cattle, horses, donkeys, camels, goats, sheep, chickens or other poultry, or beehives.
<sup>3</sup> Indicators from Nyangatom are based on the number of live births (including those children still living as well as those who subsequently died) and number of miscarriages
<sup>4</sup> The value in the DHS 2016 column corresponds to the mean number of live births in SNNPR for women who have given birth in the last 5 years. Compared to indicators of fertility from EDHS 2016, the Total Fertility Rate (TFR), calculated for the 3 years preceding the survey, was 4.6 (2.3 in urban areas, 5.2 in rural areas, and 4.4 in SNNPR).
<sup>5</sup> Estimated as the difference between the total number of children ever born alive and the number of (male and female) children who ever died. The value in the DHS 2016 column corresponds to the mean number of living children in SNNPR for women who have given birth in the last 5 years.
<sup>6</sup> p-value of the Adjusted Wald test (that accounts for the survey design) for the null Hypothesis $H_0$ : $\text{Prop}_{\text{DHS2016}} = \text{Prop}_{\text{Nyangatom2017}}$ , with a level of significance of 5% ['Prop' stands for 'mean' or 'proportion'].

**Table 2.** Problems accessing health care (Nyangatom, 2017)

|  | n | Weighted % | SE<sup>1</sup> |
| --- | --- | --- | --- |
| Number of respondents (women) | 347 |  |  |
| Asked permission (Yes) | 307 | 91.3 | 2.2 |
| Permission granted (Yes) | 300 | 97.7 | 0.7 |
| **Barriers to care identified by respondents** |  |  |  |
| Obtaining permission to go to the doctor | 11 | 2.9 | 1.4 |
| Lack of money required for advice or treatment | 100 | 25.5 | 9.6 |
| Distance to health facility | 147 | 43.3 | 7.5 |
| Not wanting to go alone | 148 | 44.0 | 4.7 |
| Other<sup>2</sup> | 41 | 11.8 | 0.5 |

**Source:** Nyangatom sample
<sup>1</sup> Linearized standard error.
<sup>2</sup> Of the 11.7% (41/347) of women responding that “Other” factors prevented them from seeking health care, the majority (69.7%, 28/41) reported that this problem was due to the requirements of caring for livestock and domestic chores.

### Problems Accessing Health Care

Women reported that the primary impediments to receiving health care were not wanting to travel alone (44.0%, 95% CI: 25.6-64.2%) and distance to the health facility (43.3%, 95% CI: 17.0-74.0%). Compared to regional data from the 2016 EDHS in which 76.7% of women reported being responsible for decisions about her own health care, only 54% percent of Nyangatom women in our sample reported being involved in decision-making, while 45% cited their husbands as being the sole party responsible for making decisions regarding the health care for the woman herself (Figure 2). However, only a small proportion (2.9%, 95% CI: 0.4-20.0, 11/347) of women reported that obtaining permission from her husband to visit a health facility was a barrier to accessing care. Nearly all women requested permission from their husbands before traveling to a health facility (91.5%, 95% CI: 78.0-97.1%), but of these women, 97.7% (95% CI: 91.6-99.4%) reported that their husbands granted them permission all of the time (Table 2).

**Figure 2.**
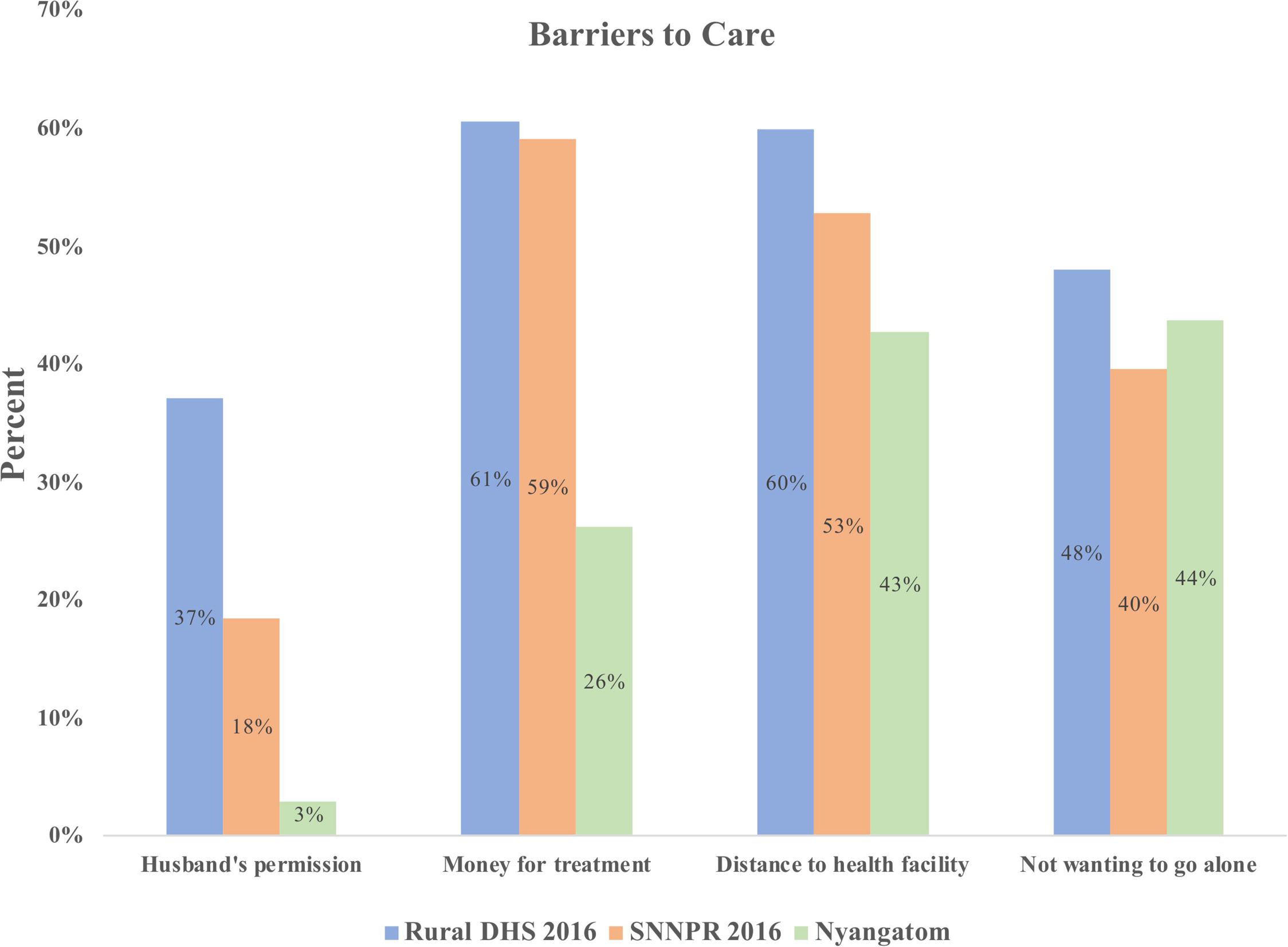
Women’s participation in decision making about their own health^1^ **Source:** Nyangatom sample and EDHS (2016). ^1^ Excluding the 1% of respondents who refused.

Women reported having more agency in decision-making for the healthcare of their children than in their own healthcare: 61.4% of women reported being involved in decision-making for the healthcare of her children. Numerous women reported that they consulted their husbands only in order to obtain money when they themselves did not possess money for transportation to the clinic or services at the clinic. A lack of alternative arrangements to care for livestock and domestic labor was cited by respondents as a major barrier to care (Table 2).

### Children’s Health

We collected health and anthropometric information from 826 of the under-15-year-old children in our sample (see SI Methods). Our final analytic sample was comprised of 547 under-five-year-old children, including 262 male children and 285 female children (Table 3).

**Table 3.** Sample size and demographic characteristics of 547 Nyangatom under-five-year-old children included in our analysis

|  | n | Month, and year of birth known (OS)<br>Weighted % | Month of birth estimated by multiple imputation (MI) |  |
| --- | --- | --- | --- | --- |
|  |  |  | Weighted % | 95% CI |
| Number of children in sample | 826 |  |  |  |
| Under-5 children | 547 |  |  |  |
| Child sex (under-5) |  |  |  |  |
| Male | 262 | 47.5 |  |  |
| Female | 285 | 52.5 |  |  |
| Total | 547 | 100.0 |  |  |
| Sex ratio (male/female x 100) | 90.4 |  |  |  |
| Child age<sup>1</sup> |  |  |  |  |
| 0 | 82 | 15.6 | 16.6 | (13.2, 19.9) |
| 1 | 97 | 17.5 | 17.8 | (14.3, 21.3) |
| 2 | 74 | 14.2 | 22.1 | (18.2, 26.0) |
| 3 | 166 | 29.1 | 21.6 | (17.7, 25.4) |
| 4 | 128 | 23.6 | 22.0 | (18.1, 25.8) |
| Total | 547 | 100.0 | 100.0 |  |

**Source:** Nyangatom sample
<sup>1</sup> Values and percentages in (OS) are based on reported month and year of birth. Percentages and 95% CI in (MI) were calculated using 120 imputed values on month of birth.

### Vaccination Status

62.8% (95% CI: 23.0-90.5%, n = 91) of the youngest children in our sample between 12-35 months of age had ever received at least one vaccination, compared to 84.1% in SNNPR. Only 14.6% (95% CI: 1.8-61.5%, 14/92) of mothers reported ever having a vaccination card in their possession for their children ages 12-23 months compared to 42.3% of children in this age range regionally in the 2016 EDHS, while among children ages 24-35 months, 8.8% (95% CI: 0.6-61.0%, 5/53) of mothers reported ever having been issued a vaccination card, versus 34.7% in the 2016 EDHS. An even smaller percentage (3.9%, 6/145) of respondents were able to present their vaccination card to researchers (6.4% for Nyangatom children ages 12-23 months compared to 28.8% of the corresponding age group in the region as reported by the 2016 EDHS). Only 5.9% (95% CI: 0.2-62.5%, 5/91) of mothers reported knowing the purpose of the vaccinations her child had received (SI Table 3).

### Treatment for Symptoms of Infectious Disease in Children

Nyangatom women in our sample sought health treatment for their children under five years of age at rates similar to national samples. In cases of diarrheal illness, 49.6% (95% CI, 32.7-66.4) of Nyangatom mothers whose children had diarrhea in the past two weeks sought treatment compared to 47.8% regionally. 46.6% of Nyangatom mothers whose children had febrile illness in the two weeks preceding the survey date sought treatment for their child’s sickness, compared to 36.7% regionally. Of mothers who sought treatment for their child’s illness, the majority visited a health clinic (65.2% for diarrheal illness and 62.4% for febrile illness) rather than a traditional provider (32.2% for diarrheal illness and 32.6% for febrile illness). See SI Table 4 for complete summary statistics on treatment-seeking.

### Anthropometric Data

Estimates of HAZ/stunting, WAZ/underweight, and WHZ/wasting were based on samples comprising 339, 190, and 182 under-five-year-old children, respectively (see SI Methods). Mid-upper arm circumference (MUAC) measurements were recorded for 339 children aged 6-59 months. Nutritional status was favorable among Nyangatom children in comparison to rural data from the 2016 EDHS. This finding was most significant with respect to WHZ/Wasting. 13.3% of under-five-year-old children in our sample met criteria for wasting compared to 24.8% from the 2016 rural EDHS, and only 1.1% met criteria for severe wasting compared to 7.3% of children in the 2016 rural EDHS (Figure 3). The lower prevalence of stunting and wasting observed among our sample as compared to the 2016 DHS calculated using height and weight measurements was consistent with our finding that very few children (2.1%) met the criteria for severe acute malnutrition (SAM), calculated using MUAC. See Table 4 for complete results on stunting, wasting, and other anthropometric data.

**Figure 3.**
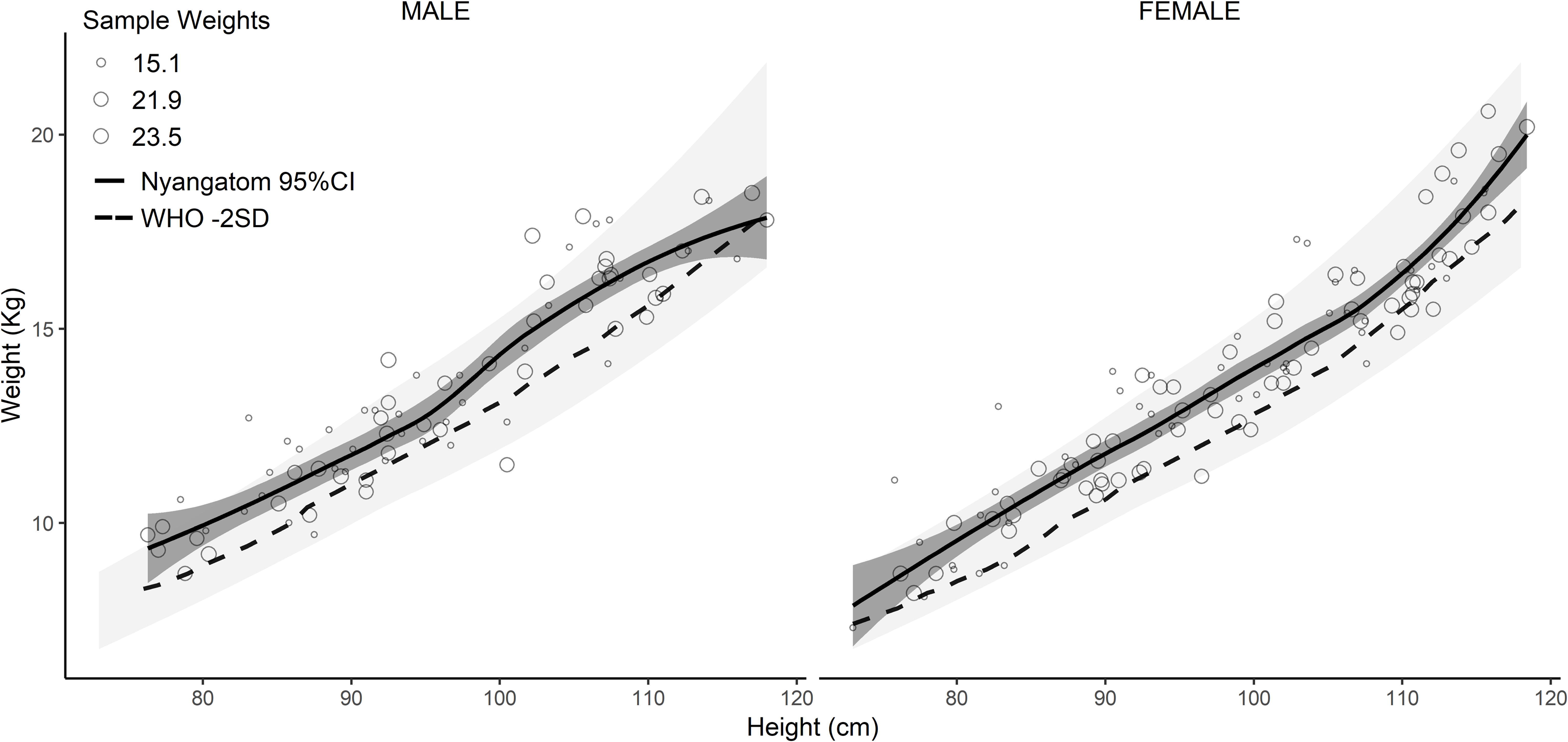
Severe Acute Malnutrition and Wasting by sex based on Weight-for-Height (WHZ) of Nyangatom under-five-year-old children in comparison to WHO Height-for-Weight reference chart values. The lightly shaded region indicates the WHO median to 3 SDs, while the darker shaded region is the 95% CI of the Nyangatom sample. Children falling below the dashed line (<-2SD) meet the WHO definition for wasting, while those with HW <-3SD meet criteria for SAM. **Source:** Nyangatom sample and data from WHO-UNICEF 2009^16^. Note: Sample restricted to children with valid measures for WHZ (using the original sample, n=73). Observation symbol size represents the probability proportional to size sample weighting (15.1, 21.9, and 23.5) used in the Loess smoothed regression line and 95% CI. Where month of birth was not known we used multiple imputation to estimate date of birth (month/year; see SI methods).

**Table 4.** Nutrition status of under-5 children, Nyangatom 2017

| Anthropometric failure | Ethiopia DHS 2016 (Weighted Mean/%) |  | Month, and year of birth known |  |  | (1) vs. (2) p-value<sup>1</sup> | Month of birth estimated by multiple imputation |  |  |
| --- | --- | --- | --- | --- | --- | --- | --- | --- | --- |
|  | Rural | SNNPR (1) | n | Weighted (Mean/%) (2) | 95% CI |  | n | Weighted (Mean/%) (3) | 95% CI<sup>b</sup> |
| HAZ / Stunting |  |  |  |  |  |  |  |  |  |
| Sample size (n) |  |  | 164 |  |  |  | 337<sup>a</sup> |  |  |
| Mean HAZ (95% CI) | -1.5 | -1.5 |  | -0.88 | (-1.77, 0.01) | 0.10 |  | -0.83 | (-1.09, -0.56) |
| Stunted n (%) | 39.9 | 38.6 | 59 | 35.3 | (23.9, 48.8) | 0.38 |  | 33.7 | (28.1, 39.3) |
| Severely stunted | 18.4 | 20.2 | 36 | 21.2 | (11.5, 35.8) | 0.76 |  | 19.4 | (14.7, 24.1) |
| WAZ / Underweight |  |  |  |  |  |  |  |  |  |
| Sample size (n) |  |  | 78 |  |  |  | 190 |  |  |
| Mean WAZ (95% CI) | -0.5 | -0.2 |  | -0.89 | (-1.76, -0.03) | 0.07 |  | -0.77 | (-0.96, -0.58) |
| Underweight n (%) | 10.1 | 6.0 | 21 | 26.9 | (17.4, 39.3) | 0.01 |  | 18.5 | (12.4, 24.6) |
| Severely underweight | 3.0 | 1.7 | 6 | 7.1 | (1.0, 36.4) | 0.22 |  | 4.5 | (1.4, 7.6) |
| WHZ / Wasting |  |  |  |  |  |  |  |  |  |
| Sample size (n) |  |  | 73 |  |  |  | 182 |  |  |
| Mean WHZ (95% CI) | -1.2 | -1.0 |  | -1.08 | (-1.52, -0.65) | 0.50 |  | -1.09 | (-1.21, -0.96) |
| Wasting n (%) | 24.8 | 21.1 | 14 | 18.3 | (5.2, 47.9) | 0.63 |  | 13.3 | (8.3, 18.4) |
| Severe wasting | 7.3 | 6.4 | 2 | 2.7 | (0.4, 16.6) | 0.09 |  | 1.1 | (-0.4, 2.5) |
| Mid-upper arm circumference (MUAC), cm |  |  |  |  |  |  |  |  |  |
| Sample size (Number of under-5 children >6m) |  |  | 157 |  |  |  | 336 |  |  |
| Mean (cm) |  |  |  | 13.5 | (12.8, 14.2) |  |  | 13.6<sup>a</sup> | (13.5, 13.7) |
| Severe acute malnutrition (SAM: MUAC <11.5cm) |  |  | 4 | 1.1 | (0.9, 1.3) |  |  | 2.2 | (0.6, 3.8) |

**Source:** Nyangatom sample
<sup>1</sup> p-value of the Adjusted Wald test (that accounts for the survey design) for the null Hypothesis H<sub>0</sub>: Prop<sup>DHS2016</sup> = Prop<sup>Nyangatom2017</sup>, with a level of significance of 5% ['Prop' stands for 'mean' or 'proportion'].
<sup>a</sup> Estimation sample varies across imputations; Sample sizes vary between 337-343 for HAZ/stunting, and between 336-343 for MUAC.
<sup>b</sup> Accounts for the variation in the imputed samples for age in months.
Values and percentages in (OS) are based on reported month and year of birth. Percentages and 95% CI in (MI) were calculated using 120 imputed values on month of birth.

## Discussion

Our results demonstrate that geospatial sampling frameworks can be used to construct representative samples of nomadic populations in a logistically feasible, cost-sensitive manner. Our approach was designed to overcome the limitations of census-based sampling frames such as those used by household surveys including the DHS, the World Bank’s Living Standards Measurement Study (LSMS), and UNICEF’s Multiple Indicator Cluster Surveys (MICS). Conventional methods fail to account for population mobility, contributing to the exclusion of nomadic groups in survey data.

While enumerations using geospatial imagery have been reported at the single settlement level, ^12^ this is to our knowledge the first time a geospatial sampling frame was implemented to conduct a population-level assessment. Previous studies found that dispatching survey personnel without first determining the inhabitance status of randomized geographic regions was logistically infeasible.^8^ By enumerating currently inhabited settlements in advance, we were able to assess active population distributions to generate density estimates in a remote, difficult-to-access area. We then demonstrated the logistical feasibility of this method through our field validation.

The use of recent satellite imagery played a crucial role by providing nearly real-time information about the distribution of the target population. Of the 25 settlements selected for survey, two exhibited signs of very recent abandonment (both were active when scouted in the week prior to the intended date of survey, but abandoned by the time of survey), two showed signs abandonment at a more distant time point (demonstrating the limitations of identifying settlement status from space), and one site was not located due to a suspected faulty coordinate pair. Refining our methods in the future, we expect that we will be able to compress the timeline from satellite imagery capture to survey deployment, reducing the number of settlements abandoned by the time of the field team’s arrival. For subsequent surveys, we also recommend visual confirmation and coordinate checks for all selected settlements once the sampling frame has been generated.

Once the satellite imagery for our study was prepared, manual reconnaissance required ∼6 hours to visually enumerate settlements in an area of over 5,000 km^2^. This approach is minimally resource-intensive over large areas of terrain characteristic of pastoralist environments. Our method depends upon environmental features such as sparse vegetation and the distinctive visual signature of settlements. Though implementation may be more difficult in heavily forested settings, nomadic populations tend to inhabit areas of sparse vegetation similar to the conditions of our study area. For this reason, we believe that our methods should be generalizable to nomadic pastoralists in general, and in Sub-Saharan Africa in particular. Additionally, we believe that machine automation of parts of the reconnaissance process will be possible, reducing human intervention to verification.^17^ The increasing availability of high-resolution, high-cadence satellite imagery and software pipelines for automated pre-processing and feature extraction will reduce human reconnaissance time and increase the capacity for greater geographic coverage.

Integration with existing DHS methodology is crucial. This could be done by working with Ministries of Health and other national partners to identify pastoralist regions in which a geospatial approach should be implemented in favor of traditional census-based enumerations. Survey weighting and population estimates remain a challenge. However, we feel that our approach provides the possibility of more accurate population estimates than existing methods.

Compared to the 2016 EDHS sample regionally and in rural areas nationally, Nyangatom women demonstrate a relatively high uptake of ANC services. Recent NGO- and government-supported initiatives including the provision of free MCH services at the nearest health clinic as well as recent road construction increasing access to transportation may explain this trend. Our finding that, under certain conditions, pastoralist women seek treatment at health facilities at rates comparable to regional non-pastoralist populations counters narratives that pastoralists tend to underutilize health services because of cultural beliefs or preferences for traditional healers.^18^ At a minimum, our findings suggest that such reports cannot be extrapolated to characterize the health-seeking behavior of pastoralist populations in general.

Despite higher ANC-seeking rates, skilled birth attendance (SBA) was significantly lower (6.4%) among the Nyangatom compared not only to the average from rural areas nationally (21.2%), but also lower than the lowest regional figure reported by the 2016 Ethiopia DHS (16.4% in the Afar region). The national trend in the reduction of at-home births has not reached this population, and the vast majority of births occur at home (91.2%) compared to the national rural average (79.0). Only 6.8% of women in our sample delivered at a health facility vs. 19.6% in rural areas nationally. The determinates of this discrepancy would benefit from further research. ANC services may be more accessible for pastoralist women since they can time their visits when their encampments are settled in a location favoring travel to a health facility. By contrast, pastoralist women may be unable to time their deliveries to give birth at a health clinic, as they frequently continue to migrate with mobile livestock encampments and perform domestic labor in locations distant from health facilities throughout their pregnancy. Though the generalizability of this finding to other pastoralist populations should be investigated, this finding highlights opportunities to target programming, outreach, and research towards SBA and delivery practices among pastoralists.

Though limited by lack of reliable documentation, Nyangatom children in our sample had low vaccination coverage compared to the EDHS, with only 62.8% of Nyangatom children ages 12-35 months having ever received a vaccination versus 84.1% of children regionally in SNNPR. The majority of our respondents did not retain possession of their vaccination cards even when these documents had been provided to them, or produced documents that were unreadable due to weathering from dirt and mold. As a result, mothers were not able to furnish definitive documentation of full immunization coverage for any of the children in our sample. By mothers’ recall, these findings are consistent with other reports of low rates of vaccination coverage among other pastoralist populations.^19–21^

Ethiopia’s National Expanded Programme on Immunization has identified pastoralists as the predominant population at risk for missing immunization services.^22^ Many respondents in our sample reported that their children missed vaccination events while residing in a different area than where coverage was provided. These accounts are similar to those of Fulani pastoralists in Nigeria, the majority of whom reported they were missed by supplementary immunization activities during an oral polio vaccination campaign due to absence of the child at the time of health workers’ visit.^23^ Such studies provide support for the argument that sampling bias may lead to substantial inaccuracies in estimates of vaccination coverage,^24^ and suggest that even mobile vaccination programs may be insufficient if they are not informed by knowledge of seasonal transhumance patterns.

Our findings suggest that conventional narratives of factors preventing pastoralist women from seeking health care may be inaccurate in some settings. Among our study population, participants reported that money, women’s empowerment, and distance to a health facility were not as significant an impediment to seeking care as insecurity and the issue of domestic labor. Subsistence livelihoods rely on daily labor from female household members such as milking and watering livestock, and collecting firewood and water. While for urban populations lack of access to money may be a major impediment, among pastoralists, the requirements of household labor may be more significant. Likewise, living in a patriarchal society such as that of the Nyangatom cannot be interpreted to mean that obtaining permission from husbands is a significant barrier to care. This finding differs from previous studies reporting that men’s control over health care decisions is a primary barrier to care for pastoralist women.^21, 25^ Although most women in our sample needed to obtain their husband’s permission to seek care, the vast majority (97%) reported they were granted this permission all of the time, and that this was not therefore a barrier to care.

Nyangatom children in our sample had a lower prevalence of severe acute malnutrition (SAM) and stunting compared to national averages. Though we believe that these results represent adequate nutritional status of the pastoralist children in our sample, the issue of survivorship remains a possibility if undernourished children do not survive to appear in our sample. For children above approximately 6 months, our results are consistent with findings from previous research demonstrating that settled children may have poorer nutritional status. This has been attributed to decreased access to milk and animal products associated with a sedentarized diet, even when total caloric intake may be greater.^26, 27^ There may be variability between populations, as other studies have found children of nomadic families to have unfavorable nutritional status compared to sedentary counterparts.^20, 28^ These findings underscore that arguments for sedentarization premised on improvements in childhood nutritional status (historically put forward by development agencies, national governments, and non-governmental organizations) must have an empiric basis.^27^ An unexpected finding of our study concerned the misuse of veterinary medicine for self-treatment. Numerous mothers reported administering livestock medicine to their children, including the use of injectable cattle ivermectin applied directly to the eyes for pediatric ophthalmic illness, and veterinary oxytetracycline as an orally administered antitussive. This potentially concerning practice merits further research.

The limitations of this study highlight several challenges inherent in surveying pastoralist populations. First, most Nyangatom women in our study population were unable to accurately date events and ages. As a result, it was necessary to impute ages for a number of children in our sample, with a potential impact on indicators depending on children’s precise age. Second, the domestic labor schedules of our participants constrained both the hours during which it was feasible to collect data as well as participants’ susceptibility to survey fatigue, limiting the length of our survey. During data collection, women were occupied chasing birds away from their fields and often left their dwellings before dawn. For similar reasons, we were unable to interview household heads, who herded cattle during the day and were rarely present in the same location as women participants.

Our findings suggest several ways in which MCH interventions among pastoralists can be improved. First, vaccination outreach campaigns should coordinate their timing to occur when settlements are at their largest size (among the Nyangatom this is during the wet season), in order to avoid missing inhabitants of livestock encampments dispersed across remote grazing territory in times of scarcity^26^. Second, alternative record-keeping systems for immunizations should be developed and implemented with pastoralists. Nomadic households are unlikely to retain these documents, and paper cards will not weather exposure to the elements incurred during frequent migrations.

Additionally, our findings suggest that men’s control over health care decisions may not be a major barrier to pastoralist women’s ability to seek care even in cases where a husband’s permission is required. By contrast, domestic labor and responsibilities such as caring for the family’s livestock, homestead, and other children appear to be significant and under-recognized barriers to pastoralist women seeking care. Efforts targeting these domains may be more effective in improving MCH among pastoralists, and empirically informed studies investigating the determinants of health-seeking behavior should be used to guide programming and decision-making. Interventions focusing on men’s awareness, for example, may not be as effective or impactful as providing secure, expedient transportation to a clinic to minimize time away from the household.

Despite pastoralist groups’ long history of invisibility in population data, our sampling design and successful field deployment indicate that geospatially derived sampling frames hold potential for the systematic inclusion of mobile populations in household surveys. This method is a viable alternative for use in pastoralist regions, and has the potential to increase the representation of mobile populations in survey data. The disparities we found between the study population and DHS-derived country estimates for certain MCH indicators suggest that without the implementation of sampling strategies capable of accounting for mobility, key indicators in data used to monitor progress on Sustainable Development Goals and other targets may present an inaccurate picture of pastoralists’ health status.

Above all, we propose that policymakers and national governments implement the use of geospatial sampling frames in pastoralist regions to reduce under-coverage and prevent bias in national estimates. This is a crucial step towards designing health systems and implementing longitudinal health surveillance for underserved mobile groups. Such an approach can aid in more effectively targeting the limited resources of national Ministries of Health, and serve as an overdue foundation to include mobile pastoralists in frameworks for achieving universal health coverage.

## Acknowledgements

We thank the College of Development Studies at Addis Ababa University for hosting our research, including Drs. Dilu Shaleka and Emezat Mengesha; Dr. Elias Alemu and Dr. Hannah Getachew of Hawassa University School of Behavioral Sciences; Gemechu Kuffa and colleagues at the SNNPR Health Bureau; and the Nyangatom Administration, especially Lore Kakuta. We are particularly grateful for logistical support provided by Semie Anbesse, as well as from Zelalem Worku. We thank Solomon Tilahun for providing shapefiles used in geospatial analysis, and Mina Mousa of Stanford University School of Medicine’s Information Resources and Technology for remote technical support which was essential for our fieldwork.

