## Supplemental Information for "Making Pastoralists Count: Geospatial Methods for the Health Surveillance of Nomadic Populations"

### SI Methods

Calculation of Study Area:

Nyangatom study area calculated using *woreda* boundaries accessed from: United Nations Office for the Coordination of Humanitarian Affairs office in Ethiopia: https://data.humdata.org/dataset/ethiopia-population-statistics

Analysis

Means, proportions, rates, and prevalence of health indicators were calculated using Stata software version 14.2 (StataCorp, College Station, TX) and anthropometric scores were calculated using the command ‘zscore06’ available in Stata.[1]

Sample size calculation

We followed a standard and well-established procedure for the sample size calculation of the number of women to be interviewed for this study. We considered the proportion of stunted children (r), a critical indicator of the undernutrition status of the under-age-five population, as the reference indicator for our calculations. We used the following formula for this purpose:[2]

$$n = \frac{(z_\alpha)^2 \cdot r \cdot (1-r) \cdot DEFF \cdot k}{p \cdot n_h \cdot e^2},$$

where $n$ is the number of women to be selected in the sample; $z_\alpha^2$ is the statistic from a standard normal distribution with a level of significance of $\alpha$; $r$ is an estimate of a key indicator to be measured by the survey; DEFF is the design effect that adjusts the variance estimation resulted from a complex survey design in comparison with that obtain using a simple random sample design (more details below); $k$ is an adjusting factor that accounts for an anticipated non-response rate; $p$ was the proportion of the target population relative to the total population of reference upon which $r$ is based; $n_h$ is the average number of household members; and $e$ represents the standard error to be attained. A selection of these parameters was conducted information from comparable sources and considering a balanced approach in which we could attain the maximum possible sample (with a conservative assumption of maximum variation) and a reasonable statistical power (>70% or 80%), but also accounting for the limited financial resources available to conduct the study.

A sample size of 400 women was selected, after considering a level of significance of 95% (with $z_{\alpha/2}$=1.96 that corresponds to a level of significance of $\alpha$ = 0.05); a stunting prevalence of 50% ($r=0.5$) (which provides the maximum variance for a proportion estimate) after observing a prevalence of stunting of 39% in SNNPR, and similar or even higher in other regions (see 'Statistical power' section below); a design effect DEFF=2.0 (DEFF values of the DHS 2016 survey for the SNNPR region were 1.86, 1.09, and 1.99 for HAZ, WHZ, and WAZ, respectively); an anticipated non-response rate of 10% or $k=1.1$ (the response rate in SNNPR was 95.4% in DHS 2016); a proportion of under-age-five children of $p=15\%$, estimated from population values from UNICEF Ethiopia from 2012 (https://www.unicef.org/infobycountry/ethiopia_statistics.html); an average household size of $n_h=6$ members per household (a slightly higher size than the average of 4.9 in rural areas); and a margin of error of $e=7.5\%$, obtained as the 15% of our reference prevalence value of $r=50\%$.

Statistical power

To assess the statistical power that we expected to attain for the hypothesis testing of a stunting prevalence of 50% in Nyangatom, we simulated the power under different alternative proportions observed in distinct regions of Ethiopia in 2016 (EDHS 201); namely: Tigray 39%, Dire Dawa 40%, Affar 41%, and Benishangul-Gumuz 43%. In Supplementary Figure 1(a), we observe that had the true value of the stunting prevalence in Nyangatom had been closer to 40%, we would obtain a statistical power around or higher than 90% with sample sizes above 300 women (assuming we could measure at least one under-age-five child per mother). That is, with a sample size of 400 children, our power would have been larger than 95%. During the survey, however, we were only able to interview only 347 women with 547 under-age-five children, but only for a subsample of 164 of those children we were able to obtain reliable estimates of stunting. With this small sample, we estimated again the power that we effectively obtained for our initial hypothesis of 50% of stunting, but now using the alternative values that we were able to obtain in our study (35% with our original sample, and 34% with the imputed sample -see below). The results of this exercise are illustrated in Supplementary Figure 1 (bottom panel), where we can observe that we were able to obtain a statistical power above 95%, even with this reduced sample.

<u>Sampling weights</u>

The Nyangatom sample was selected in two stages. In the first stage, 3 primary sampling units (PSUs) were selected with probability proportional to sampling size from a list of 15 PSUs that covered the entire sampling area. In the second stage, 342 mothers of reproductive age in the selected units (clusters) were interviewed on maternal and child health. We constructed sampling weights at the dwelling and child levels to ensure representativeness of the survey results for the Nyangatom region. Sampling weights were constructed as follows:

If $P_{1i}$ and $P_{2i}$ represent sampling probabilities for the first- and second-stages in the cluster $i^{th}$ (respectively), $a$ is the number of PSUs selected, $M_i$ the number of huts (dwellings) in the $i^{th}$ cluster, obtained from a counting of huts from satellite imagery, and $\sum M_i$ the total number of huts in the region, then the probability of selecting the $i^{th}$ PSU is given by $P_{1i} = \frac{aM_i}{\sum M_i}$.

In the second stage, the selection probability for each household in the cluster $i$ was calculated as: $P_{2i} = \frac{g_i}{L_i}$, where $g_i$ represents the number of huts selected in the cluster $i$, and $L_i$ the total number of huts in the corresponding PSU. As a complete listing of huts in the selected clusters does not exist and conducting a manual count of all huts in the area was unfeasible, we estimated $L_i$ using a count of huts from satellite imagery in a sample of villages. We then used this to generate an estimate of average village size and extrapolated this to the cluster level.

If $c_i$ is the average number of children interviewed per mother (hut) and $si$ the average number of alive children per mother (hut) in the population (we assumed an average number of family members per hut of 6 based on information from the EDHS,[3]) then the probability of selecting one child for interview per hut in the $i^{th}$ cluster was computed as: $P_{3i} = \frac{c_i}{s_i}$.

The sampling weights for each selected hut in cluster $i$ were then estimated as the inverse of its overall selection probability: $W_{hi} = \frac{1}{P_{1i} \cdot P_{2i}}$. Similarly, the sampling weights for each selected child in the cluster was computed as: $W_{ci} = \frac{1}{P_{1i} \cdot P_{2i} \cdot P_{3i}}$.

Design Effects

The standard errors or confidence intervals for all means, proportions, rates, and prevalence of health indicators in this study considered the complex survey design (CSD) of the Nyangatom sample (two-stage cluster design). Compared to a simple random sample design (SRS), the variance of estimates under the CSD consider the proper probability selection of sampling units (as explained in the previous section). To assess the difference in the precision between estimates obtained from both designs, Kish[4] proposed an indicator commonly known as design effects (DEFF) that is calculated using the expression $DEFF = \frac{V_{CSD}(\hat{y})}{V_{SRS}(\hat{y})}$, where $V_{CSD}(\hat{y})$ and $V_{SRS}(\hat{y})$ represent the variance of the estimator $\hat{y}$ obtained under *CSD* and *SRS* survey design schemes, respectively. DEFF is an inflation factor that indicates how the variance increases/decreases (usually increases) when adopting a complex design relative to the SRS design. To illustrate the magnitude of the effect of our survey design strategy, we replicated the indicator of Table 2 (main manuscript) and computed standard errors without accounting for the cluster design and the corresponding DEFFs (Supplementary Table 1). For instance, the variance of the proportion of women obtaining permission to go to the doctor would be 2.3 times lower had we adopted a SRS design for the Nyangatom survey.

**Supplementary Table 1** Problems accessing health care (Nyangatom, 2017)

| | Weighted % (95% CI) | | DEFF |
|---|---|---|---|
| | SRS[1] | CSD[2] | |
| Asked permission (Yes) | 91.5 (88.1, 94.1) | 91.5 (78.0, 97.1) | 1.7 |
| Permission granted (Yes) | 97.7 (95.2, 98.9) | 97.7 (91.6, 99.4) | 0.7 |
| Significant barriers to care | | | |
| Obtaining permission to go to the doctor | 2.9 (1.6, 5.2) | 2.9 (0.4, 20.0) | 2.3 |
| Lack of money required for advice or treatment | 25.5 (21.2, 30.4) | 25.5 (3.7, 75.1) | 16.4 |
| Distance to health facility | 43.3 (37.9, 48.8) | 43.3 (17.0, 74.0) | 7.7 |
| Not wanting to go alone | 44.0 (38.7, 49.5) | 44.0 (25.6, 64.2) | 3.1 |
| Other[1] | 11.7 (8.7, 15.7) | 11.7 (9.6, 14.3) | 0.1 |

**Source:** Nyangatom sample

[1] Sampling weights were included to obtain unbiased estimates.
[2] CIs were estimated using linearized standard errors.

Anthropometry

For anthropometric analysis, additional exclusion criteria were adopted on the basis of missing values of height (193 children excluded) or weight (357) measurements, or implausible values that resulted in the exclusion of an average of 21 additional children (average over 120 imputed samples of age in months: 14 (3.9%) children with implausible values for HAZ were discarded in every imputed sample in average, 0 (0%) for WAZ, and 7 (3.7%), more details of the imputation procedure below) whose height-for-age z score (HAZ) was less than -6 or greater than 6, weight-for-age z score (WAZ) was less than -6 or greater than 5, or weight-for-height z score (WHZ) was less than -5 or greater than 5, according to the World Health Organization child growth standards.[5] Mothers in our sample frequently declined to allow researchers to measure babies under 6 months of age; therefore, the left-most tail of our anthropometric curve is sparsely populated.

Multiple Imputation (MI) and exclusion criteria based on the variable age

In this study, we conducted analysis for under-five-age children, but age in months is an essential input for the assessment of nutritional status. However, information at this level of detail was only reported for a subgroup of children. Among the 826 alive children in our sample, 218 (26%) had the exact date of birth (day/month/year), 328 (40%) the month/year, and 759 (92%) only the year of birth. With this information, we identified 547 (66%) under-five-age children (children born after August 2012), but 284 of them (52%) reported missing birth month. Therefore, age in months was computed for the remaining 263 children, and for the 284 children after imputing their missing month of birth. We used multiple imputation (MI), aiming to increase the sample size and statistical power of our undernutrition analysis.

*Goodness of fit of MI data*

Before the imputation, we explored the use of available information on sex, height, and weight as potential covariates to increase the accuracy of the model and to deal with the requirement that missing data are missing at random (MAR).[6] Specifically, when the data generation process is

Missing Completed at Random (MCAR) or MAR, MI provides unbiased and reliable results even with large proportions of missing data, according to as simulation study from Lee and Huber (2011).[7] In our sample, we did not appreciate a substantive difference in the distribution by age of the month of birth variable, nor a significant mean difference of weight and height between samples with and without missing month of birth. In addition, the inclusion of these covariates did not improve the goodness of fit of the MI model, and then we decided to exclude them from the final model specification. We imputed the missing entries of birth of month in the incomplete dataset 120 times (more than 2 times the percentage of missing values for this variable in the reduced sample –52%), drawn from a multinomial logistic distribution, which resulted in 120 complete datasets. Supplementary Figure 2 shows that the original month of birth variable corresponding to the first imputed sample followed similar patterns and levels of that variable in the original sample for the period between 2013 and 2017, providing evidence in favor of the MAR assumption and the robustness of the MI model (we did not show the distribution for the remaining 119 replications, but the results are similar). We also compared the distribution by age of the under-age-five sample before and after imputation (see Table 3 in the main manuscript), and compared to the corresponding distribution of Ethiopia in 2006—overall and by rural/urban area—using information from EDHS 2016. After imputation, we did not find statistically significant discrepancies in the distribution between the imputed sample and the DHS data across age groups (see overlap of CIs in Table 3 in the main manuscript). In addition, we reported 95% CIs for indicators that account for the variation of the imputed values of age in months across the 120 imputed samples of acute, chronic, and severe undernutrition indicators (Wasting, Underweight, Stunting) in Table 4 of the main manuscript, and we conducted further goodness of fit tests of the MI approach as reported in the Supplementary Table 6. These tests are based on guidelines from previous studies that have tested the reliability of MI in terms of efficiency of point estimates or estimation of standard errors. [6,8–11,7]

Early literature focused on statistical efficiency and concluded that high efficiency can be obtained even with a small number of imputations (e.g., 10 imputations is needed to get 95% effciency[9]). The relative efficiency was around 99% for all age groups in our sample, which reflects how well the distribution of age was estimated. Standard errors were calculated using Rubin's (1987)[6] formula that combines variability within and between data sets. The standard errors were then estimated using the underlying variability of the estimated proportion of under-age-five children for each age group across the multiple datasets. The results indicate a moderate inflation in the standard errors, resulting from the MI procedure that added additional uncertainty into the estimates, at all ages (with a larger % increase at age 3 of 11%), and increasing slightly relative to the original standard errors, but with a more stable pattern. The results in the table also indicate that a proportional increase in total sampling variance that is due to missing information (RVI) varied across age groups, from 7.7% for age 0 to 24% for age 3. Similar results are derived from the fraction of missing information (FMI), which indicates that 7.1% of the total sampling variance was attributable to missing data at age 0, while it contributed to 24.0% of the total at age 3.

*Anthropometric assessment using MI data*

After the imputation of the month of birth, we estimated the age in months for children with missing information for the assessment of undernutrition measures (HAZ, WAZ, WHZ, stunting, underweight, wasting, and corresponding severe measures) using the imputed complete samples. Mean or proportion values were then assessed using the 120 replicated samples, as well as 95% confidence limits of weighted estimates that accounted for the additional variation of imputed samples, but not for the cluster variance. The analytic sample for the anthropometric assessment increased after MI (to 339, 190, 182, and 339 observations in the replicated samples that were used for the estimation of HAZ/Stunting, WAZ/, WHZ/ Wasting, and SAM, respectively –see Table 4 in the main manuscript).

2 United Nations, editor. Designing household survey samples: practical guidelines. New York: United Nations, 2008.

3 Central Statistical Agency. Ethiopia Demographic and Health Survey 2016. 2016. http://dhsprogram.com/publications/publication-FR328-DHS-Final-Reports.cfm.

4 Kish L. Survey Sampling. New York: Wiley, 1965.

5 World Health Organization. WHO child growth standards and the identification of severe acute malnutrition in infants and children: joint statement by the World Health Organization and the United Nations Children's Fund. 2009. http://apps.who.int/iris/bitstream/handle/10665/44129/9789?sequence=1.

6 Rubin DB. Multiple imputation for nonresponse in surveys. New York: John Wiley & Sons, 1987.

7 Lee JH, Huber JJ. Multiple imputation with large proportions of missing data: How much is too much? Stata Users Group, 2011.

8 Graham JW, Olchowski AE, Gilreath TD. How Many Imputations are Really Needed? Some Practical Clarifications of Multiple Imputation Theory. *Prev Sci* 2007; **8**: 206–13.

9 Schafer JL. Multiple imputation: a primer. *Stat Methods Med Res* 1999; **8**: 3–15.

10 White IR, Royston P, Wood AM. Multiple imputation using chained equations: Issues and guidance for practice. *Stat Med* 2007. https://onlinelibrary.wiley.com/doi/full/10.1002/sim.4067 (accessed March 20, 2019).

11 Bodner TE. What Improves with Increased Missing Data Imputations? *Struct Equ Model Multidiscip J* 2008; **15**: 651–75.

### SI Tables

**Supplementary Table 2.** Percent distribution of women of reproductive age who received antenatal care (ANC) for the last birth by type of provider[1], number of ANC visits, selected ANC services, and place of delivery: sample size and % (95% CI) (Nyangatom, 2017)

| | Ethiopia DHS | | | | | | Nyangatom 2017[3] | | |
|---|---|---|---|---|---|---|---|---|---|
| | 2005 | 2011 | 2016 | | | | N | % | 95% CI |
| | Rural[5] | | Rural | | SNNPR[6] | | | | |
| | %[8] | % | % | p-val[7] | % | p-val[7] | | | |
| **Ever pregnant[2]** | | | | | | | 342 | 98.5 | (82.7, 99.9) |
| **Antenatal Care** | | | | | | | | | |
| Skilled provider | 23.9 | 26.4 | 58.3 | | 69.3 | 0.55 | 242 | 71.6 | (56.0, 83.3) |
| Traditional provider | 0.5 | 0.1 | 0.3 | | 0.1 | 0.02 | 11 | 3.3 | (1.8, 6.1) |
| Other | | 10.2 | 0.2 | | 0.2 | 0.29 | 3 | 1.0 | (0.1, 10.5) |
| No ANC | 75.0 | 63.0 | 41.2 | | 30.4 | 0.26 | 86 | 24.1 | (11.0, 44.9) |
| Missing | 0.1 | 0.3 | | | | | | | |
| Total | 100.0 | 100.0 | 100.0 | | 100.0 | | 342 | 100.0 | |
| **Number of ANC visits** | | | | | | | | | |
| None | 75.4 | 63.1 | 41.2 | 0.05 | 30.4 | 0.26 | 86 | 24.1 | (11.0, 44.9) |
| 1 | 4.8 | 4.8 | 4.6 | <0.01 | | | 54 | 15.7 | (11.6, 20.9) |
| 2-3 | 11.3 | 17.6 | 26.8 | 0.18 | | | 113 | 33.9 | (20.7, 50.1) |
| 4+ | 8.1 | 14.4 | 27.3 | 0.08 | | | 68 | 20.1 | (12.5, 30.7) |
| Don't know | 0.4 | 0.1 | 0.1 | <0.01 | | | 21 | 6.2 | (4.8, 8.1) |
| Total | | | | | | | 342 | 100.0 | |
| **Average No. of ANC visits** | | | | | | | 235 | 2.8 | (2.4, 3.2) |
| **Median number of months pregnant at time of first ANC visit (IQR)** | 6.0 | 5.5 | 4.9 | | | | 244 | 4.0 | (3.0,6.0) |
| **Number of months pregnant at time of first ANC visit** | | | | | | | | | |

| | | | | | | | | | |
|---|---|---|---|---|---|---|---|---|---|
| No ANC | 75.4 | 63.1 | 41.2 | <0.01 | 30.4 | 0.26 | 86 | 24.1 | (11.0, 44.9) |
| <4 | 3.9 | 7.7 | 17.0 | 0.08 | | | 80 | 22.9 | (16.3, 31.3) |
| 4-5 | 8.2 | 13.8 | 24.6 | 0.40 | | | 91 | 27.2 | (18.0, 38.8) |
| 6-7 | 8.9 | 12.2 | 14.2 | 0.20 | | | 56 | 16.7 | (11.7, 23.3) |
| 8+ | 3.1 | 2.9 | 2.6 | 0.32 | | | 17 | 5.4 | (1.0, 25.0) |
| Don't know / missing | 0.4 | 0.2 | 0.5 | 0.07 | | | 12 | 3.7 | (1.3, 10.6) |
| **Services Delivered during ANC** | Rural | Rural | Rural | | SNNPR | | 342 | | |
| Blood pressure measured | 55.7 | 67.3 | 71.7 | | 70.8 | 0.33 | 158 | 65.8 | (47.7, 80.2) |
| Urine sample | 16.2 | 29.2 | 60.3 | | 56.7 | 0.02 | 63 | 26.0 | (12.1, 47.3) |
| Blood sample | 17.3 | 44.5 | 67.6 | | 66.0 | 0.02 | 88 | 35.4 | (19.9, 54.8) |
| Tetanus injection[4] | | | | | | | 162 | 67.3 | (35.4, 88.5) |
| **Average No. of ANC services provided** | | | | | | | 342 | 1.4 | (0.9, 1.9) |
| **Place of delivery** | | | | | | | | | |
| Home | 97.0 | 95.4 | 79.0 | | 72.5 | 0.03 | 310 | 91.2 | (61.3, 98.6) |
| Health facility / Clinic | 2.3 | 4.1 | 19.6 | | 25.5 | 0.02 | 22 | 6.7 | (0.9, 36.3) |
| Other / Missing | 0.7 | 0.5 | 1.3 | | 2.0 | 0.95 | 7 | 2.0 | (0.6, 6.2) |
| Total | 100.0 | 100.0 | 100.0 | | 100.0 | | 339 | 100.0 | |
| **Person providing assistance during delivery** | | | | | | | | | |
| Nobody | 5.8 | 4.2 | 16.3 | | 22.6 | 0.27 | 65 | 18.4 | (9.3, 33.2) |
| Skilled provider | 2.6 | 4.0 | 21.2 | | 28.6 | 0.02 | 21 | 6.4 | (0.7, 41.2) |
| Traditional / Unskilled | 28.5 | 29.4 | 46.1 | | 25.9 | <0.01 | 2 | 0.6 | (0.1, 5.7) |
| Relative / Friend/ Other | 62.9 | 62.3 | 16.3 | | 22.9 | <0.01 | 251 | 74.6 | (56.3, 87.0) |
| Total | 100.0 | 100.0 | 100.0 | | 100.0 | | 339 | 100.0 | |

[1] Skilled provider for EDHS 2005, and 2011 includes doctor, nurse, and midwife. Skilled provider for EDHS 2016 includes doctor, nurse, midwife, health officer, and health extension worker. Skilled provider for the Nyangatom population in 2017 includes health officer, nurse/midwife, and community/village health worker.
[2] Estimated using the number of ever-births or pregnancies that ended in miscarriage/still-birth.
[3] Events correspond to the last birth or pregnancy. 2017 refers to the year when the survey was conducted.
[4] Single shot of tetanus given during the ANC visit.

[5] *Traditional provider* also includes care from “Other” provider in EDHS 2005.
[6] *SNNPR* stands for Southern Nations, Nationalities, and Peoples’ Region, one of the nine regional states of Ethiopia.
[7] p-value of the Adjusted Wald test (that accounts for the survey design) for the null Hypothesis $H_0$: $Prop_{DHS2016} = Prop_{Nyangatom2017}$, with a level of significance of 5% [‘Prop’ stands for ‘mean’ or ‘proportion’].
[8] % is weighted percentage.

**Supplementary Table 3.** Vaccination ever received by youngest child aged 12-35 months (Nyangatom, 2017)

| | Ethiopia DHS 2016: (%[2]) | | | n | % | 95% CI |
|---|---|---|---|---|---|---|
| | Rural | SNNPR | p-val[1] | | | |
| No. of children aged 12-35 (only includes the youngest child per woman) | | | | 145 | | |
| Age 12-23 | | | | 92 | | |
| Age 24-35 | | | | 53 | | |
| Any vaccination ever received | 82.6 | 84.1 | 0.15 | 91 | 62.8 | (23.0, 90.5) |
| Knew vaccinations were for (Yes) | | | | 5 | 5.9 | (0.2, 62.5) |
| Ever had a vaccination card (Yes) [age 12-35] | | | | 19 | 12.4 | (4.0, 32.4) |
| Children age 12-23 | 41.3 | 42.3 | 0.05 | 14 | 14.6 | (1.8, 61.5) |
| Children age 24-35 | 30.6 | 34.7 | 0.04 | 5 | 8.8 | (0.6, 61.1) |
| Vaccination card seen [age 0-59] | | | | 6 | 3.9 | (0.3, 35.8) |
| Children age 12-23 | 29.8 | 28.8 | 0.03 | 6 | 6.4 | (0.3, 57.3) |
| Children age 24-35 | 12.6 | 18.1 | | | | |

**Source:** Nyangatom sample

[1] p-value of the Adjusted Wald test (that accounts for the survey design) for the null Hypothesis $H_0$: $Prop_{DHS2016} = Prop_{Nyangatom2017}$, with a level of significance of 5% [‘Prop’ stands for ‘mean’ or ‘proportion’].

[2] % is weighted percentage.

**Supplementary Table 4**. Treatment of symptoms of infectious disease for under-5 children (Nyangatom, 2017)

| Infectious Disease | Ethiopia DHS 2016 | | | Nyangatom | | |
|---|---|---|---|---|---|---|
| | Rural (%[2]) | SNNPR | | n | % | 95% CI |
| | | % | p-val[1] | | | |
| Diarrhea | | | | 260 | 47.1 | (31.8, 62.9) |
| Percentage for whom advice or treatment was sought | 42.5 | 47.8 | 0.70 | 126 | 49.6 | (33.4, 65.8) |
| Type of facility where treatment was sought | | | | | | |
| Clinic (public or private facility) | | | | 81 | 65.2 | (23.4, 92.0) |
| Traditional Provider | | | | 42 | 32.3 | (8.1, 72.0) |
| Other / Refused | | | | 3 | 2.5 | (0.2, 23.6) |
| Services provided | | | | | | |
| Oral rehydration solution | | | | 51 | 41.3 | (29.6, 54.1) |
| Pre-packaged ORS liquid | | | | 32 | 24.3 | (7.8, 55.2) |
| Fever | | | | 366 | 67.0 | (37.4, 87.4) |
| Percentage for whom advice or treatment was sought | 31.8 | 36.7 | 0.09 | 170 | 46.6 | (32.8, 60.3) |
| Type of facility where treatment was sought | | | | | | |
| Clinic (public or private facility) | | | | 105 | 62.4 | (24.4, 89.5) |
| Traditional provider | | | | 57 | 32.6 | (5.6, 79.9) |
| Other / Refused | | | | 8 | 5.0 | (0.9, 23.4) |
| Days of illness before treatment was sought (mean) | | | | 5.4 | | |
| Any drugs taken (Yes) | | | | 206 | 57.0 | (43.3, 69.7) |
| Antimalarial/Antibiotic/Other (not herbal) | | | | 2 | 1.0 | (0.2, 5.8 |

**Source:** Nyangatom sample

[1] p-value of the Adjusted Wald test (that accounts for the survey design) for the null Hypothesis H0: $Prop_{DHS2016} = Prop_{Nyangatom2017}$, with a level of significance of 5% [‘Prop’ stands for ‘mean’ or ‘proportion’].

[2] % is weighted percentage.

**Supplementary Table 5** Goodness of fit statistics for the estimation of age based on multiple imputation of the month of birth of under-five-age children in Nyangatom

| Age group (years) | Proportion | Standard Error (SE) | % Increase in SE | Imputation variance | | | RVI[1] | FMI[2] | Relative efficiency |
|---|---|---|---|---|---|---|---|---|---|
| | | | | Within | Between | Total | | | |
| 0 | 0.166 | 0.017 | 3.8 | 0.00027 | 0.00002 | 0.00029 | 0.077 | 0.071 | 0.999 |
| 1 | 0.178 | 0.018 | 5.9 | 0.00028 | 0.00003 | 0.00031 | 0.121 | 0.108 | 0.999 |
| 2 | 0.221 | 0.020 | 8.9 | 0.00033 | 0.00006 | 0.00039 | 0.187 | 0.158 | 0.999 |
| 3 | 0.216 | 0.020 | 11.4 | 0.00031 | 0.00008 | 0.00039 | 0.240 | 0.195 | 0.998 |
| 4 | 0.220 | 0.020 | 7.9 | 0.00033 | 0.00005 | 0.00038 | 0.165 | 0.142 | 0.999 |

[1] Relative increase in variance
[2] Fraction of missing information

**Supplementary Figure 1.** Power calculation for (a) different sample sizes and alternative proportions of stunted under-age-five children from selected Ethiopian regions (SNNPR, DD, Affar, and BG), and (b) after estimating the prevalence with an effective sample size (OS) obtained in for the Nyangaton study, and after imputing the month of birth for 52% of under-age-five children (MI).

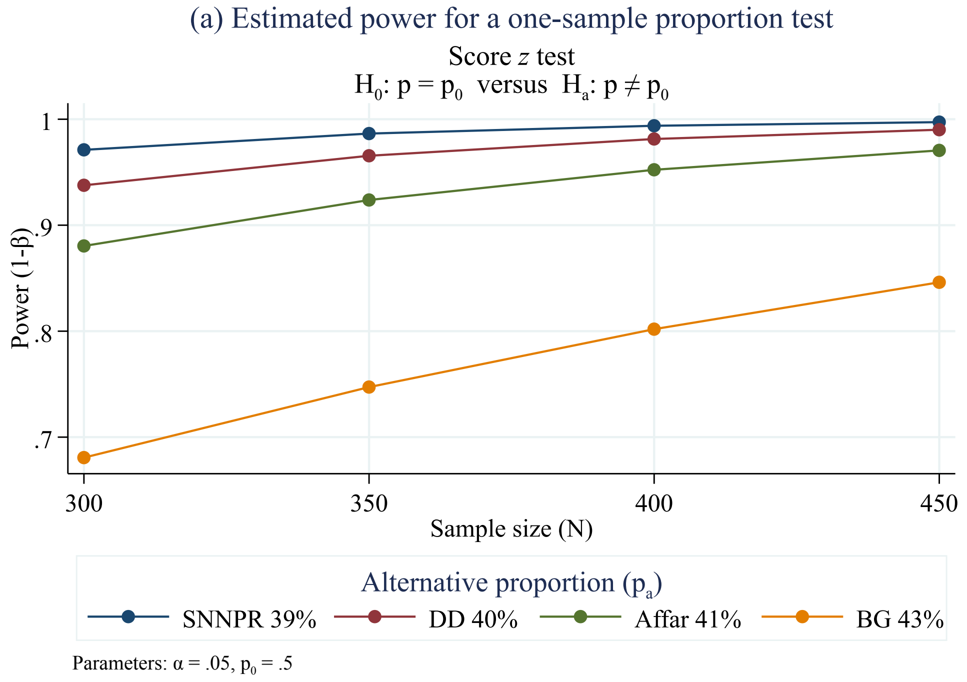


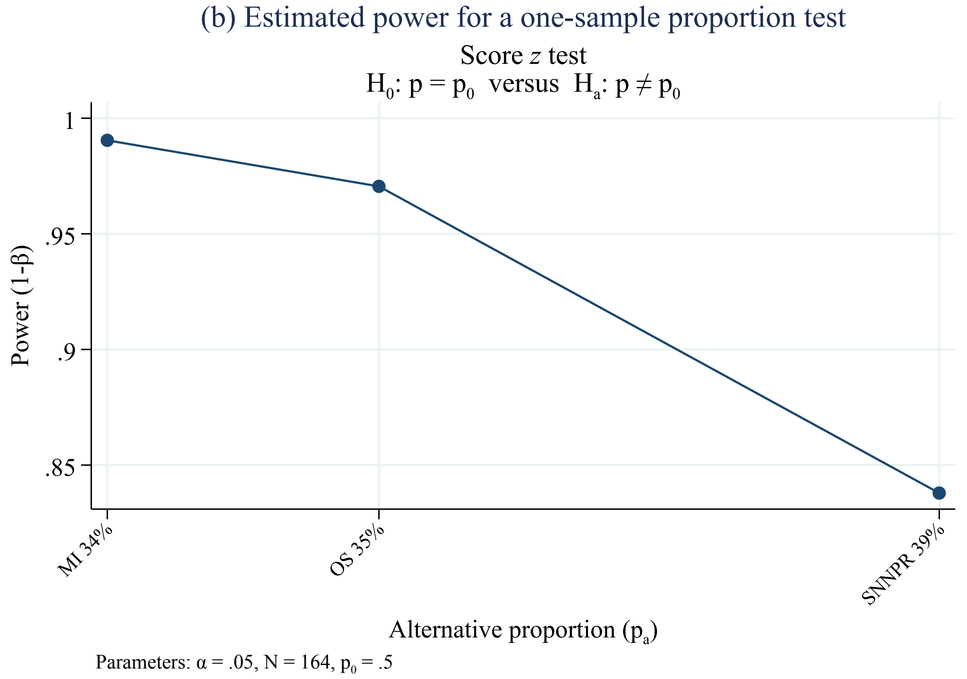


**Abbreviations:** SNNPR - Southern Nations, Nationalities, and People's Region; DD – Dire Dawa; BG - Benishangul-Gumuz; MI – Multiple Imputation; OS – Original sample

**Supplementary Figure 2.** Distribution of observed and imputed month of birth for children under-age-five from 2013-2017

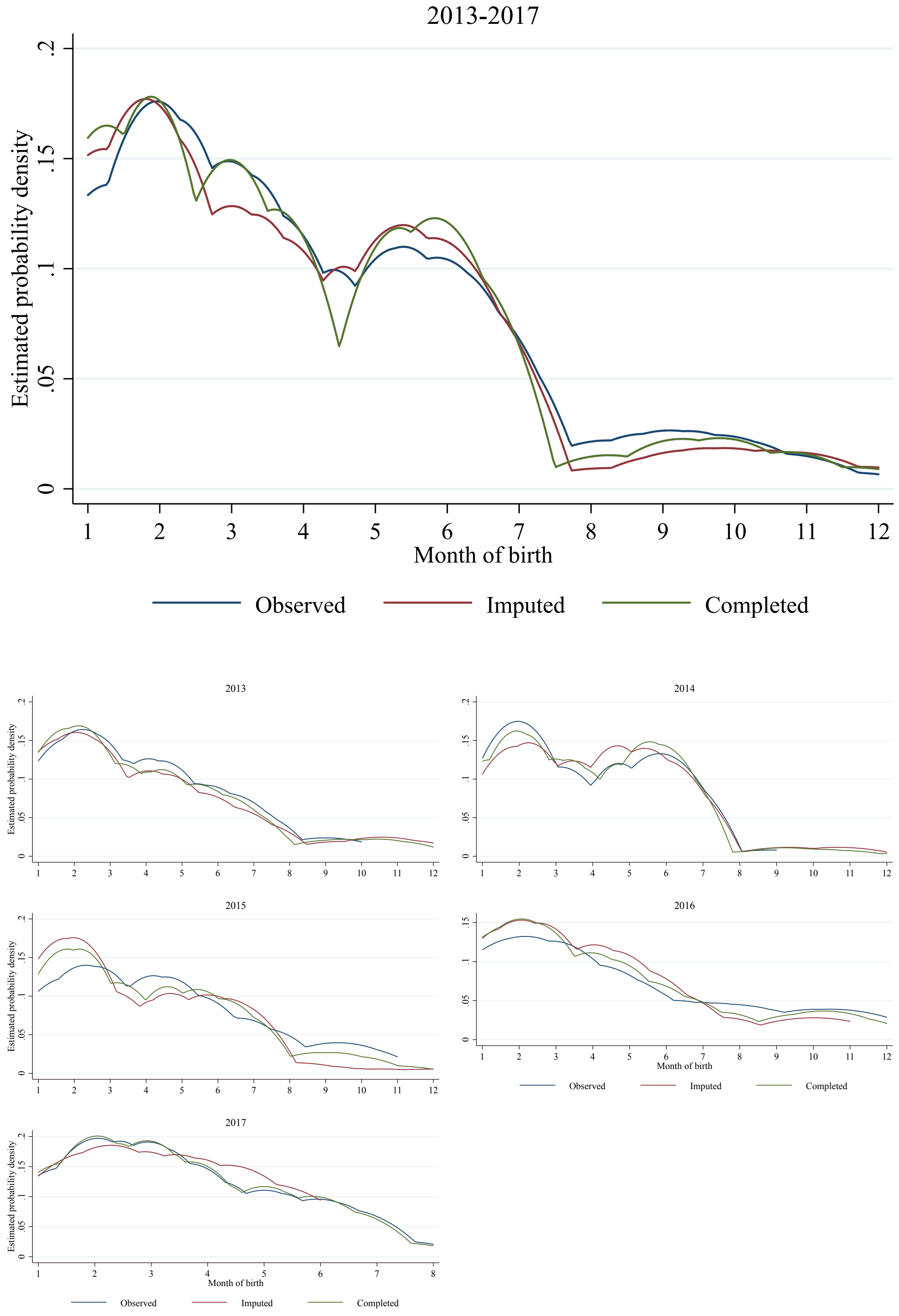


**Note:** We show the estimated probability density of the month of birth for the observed, imputed and completed (observed + imputed) samples after the implementation of the multiple imputation model. The top figure illustrates the distribution of the month of birth for the first replication of the MI model for all years combined from 2013 to 2016 (period where we identified most of under-5 children in our observed sample). The bottom figure shows the same distribution but for each year separately during the same time frame.
